# Exploring the Effect of LED-to-Photodetector Spacing on Subcutaneous Photoplethysmography for Continuous Blood Pressure Measurement

**DOI:** 10.1101/2025.09.15.676386

**Authors:** Dario Cabal, Brian Nguyen, David B. Green, Kevin Kilgore, Varun Thakkar, Michael J. Fu, Steve J.A. Majerus

## Abstract

Photoplethysmography (PPG) is widely used to measure heart rate, blood oxygenation, and more recently, blood pressure. Implanted PPG systems offer the possibility to measure similar real-time measures of cardiovascular health, however, the detection method may vary due to a lack of capillary vessels for PPG sensors to observe in muscle tissue. To improve volumetric blood detection in large muscles, without relying on the capillary density of skin, a flexible PPG sensor was developed. The sensor included multiple spacing of illuminating infrared (IR) light emitting diodes (LEDs) and a single IR photodetector. This arrangement was expected to enable detection of blood volume changes at variable distances from the sensor face, potentially at much longer depths then feasible with skin-mounted PPG devices. IR bench phantoms simulating a large blood vessel embedded in IR absorbing, tissue-mimicking rubber were developed and used to determine the sensor performance *in vitro*. A preliminary *in vivo* test used an adult rabbit to provide additional performance validation. Test results reveal an observed trend of increased SNR for deeper vessel depths for the farthest LED to detector spacing which is aligned with our initial prediction. However, ANOVA and post-hoc tests reveal that these trends did not reach statistical significance. The *in vivo* test showed a relationship consistent with relevant literature. Future experiments are required to improve the phantom’s representation of the biological setting and to confirm a reduced SNR variation for the farthest spacing.

**Clinical Relevance:** Development of a subcutaneous, continuous blood pressure sensor may provide benefits towards monitoring autonomic dysreflexia and hypertension for people affected by spinal cord injury.

## I. Introduction

Photoplethysmography (PPG) is an optical technique that has been used to measure volumetric variations of blood circulation in tissue near the presence of the sensor. In a standard reflectance-mode setup, it consists of a light emitting diode (LED) and a photodiode (PD) spaced laterally apart from one another [1]. When paired with a parallel recording of an electrocardiogram (ECG) (or with another PPG waveform taken at a point more proximal/distal from the heart compared to the first sensor) the delay between the two waveforms can be measured to calculate the pulse wave velocity. When this delay is measured from the R-wave peak of the ECG to the incoming PPG, the delay is called the pulse arrival time (PAT). When it is measured between two PPG waveforms, it is called the pulse transit time (PTT) [2]. It is well documented that a higher mean arterial pressure is inversely correlated with both PAT and PTT [3]. As such, PPG sensors potentially enable measuring blood pressure without a cuff.

PPG is usually acquired from the skin surface to noninvasively measure blood oxygenation and heart rate. Recent efforts have focused on expanding the health monitoring capabilities of implantable and subcutaneous devices by including PPG. However, mixed results have been shown when transferring this sensing technology to a subcutaneous environment [4], [5], [6], [7]. Many issues arise from motion artifacts and the inability of these sensors to consistently obtain a signal when placed subcutaneously on a muscle surface. This may occur due to variation in capillary density, or simply the low capillary density, within the tissue [6].

In this study we investigated the effect of increasing the distance between the LED and PD on PPG signal acquisition quality, which may target a wider area by passing the light through more tissue prior to reaching the PD. Light emitted from the LED must travel farther and deeper to backscatter toward the PD when the LEDs are spaced further apart. As such the light will travel through more tissue, thereby encountering more blood volume changes [8], [9]. The risk to this approach is lower optical intensity measured at the PD, which can be addressed with high-gain optical detectors in an implanted environment, where the background IR intensity is very low. Finally, by interrogating more tissue, the need to place an implanted PPG sensor in a capillary rich environment is reduced, which can potentially improve its viability for subcutaneous use.

This paper describes the construction of a PPG platform intended for implantation, which uses a variety of LED-PD spacings to enable deep tissue optical interrogation. Here, we describe initial validation of the sensor platform *in vitro* using an IR vascular phantom and *in vivo* in an acute animal study.

## II. In Vitro and In Vivo Testing methods

### A. Multi-LED PPG Sensor

The PPG sensor (Fig. 2) consisted of a flexible polyimide circuit board that housed two infrared LED with a primary wavelength of 850 nm (15414185BA210, Würth Elektronik). Reflected light induced a photocurrent on an IR photodiode with a central wavelength of 900 nm (TEMD7000X01, Vishay Semiconductor). The photocurrent was amplified by a transimpedance amplifier (OPA381AIDGKR, Texas Instruments) powered by a 3 V linear voltage regulator (NCP730ASN300T1G, On Semi). The LED output was regulated by a constant current driver from 5 mA to 100 mA (BCR430UW6-7, Diodes Incorporated).

**Figure 1.**
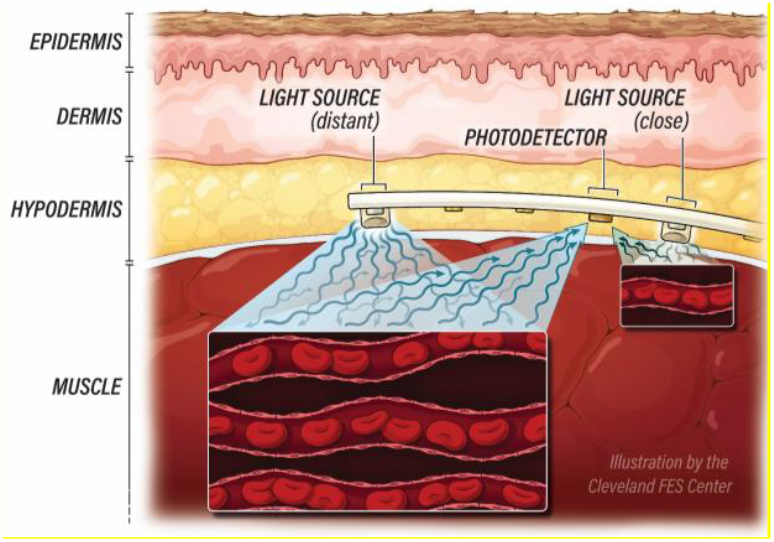
The proposed multi-LED, reflectance-mode implanted PPG system is intended to interrogate deeper tissue thickness. This increases the likelihood for optical absorbance by blood flow at multiple levels deep within tissue. This is necessitated by a lower capillary density in muscle compared to surface structures like skin.

**Figure 2.**
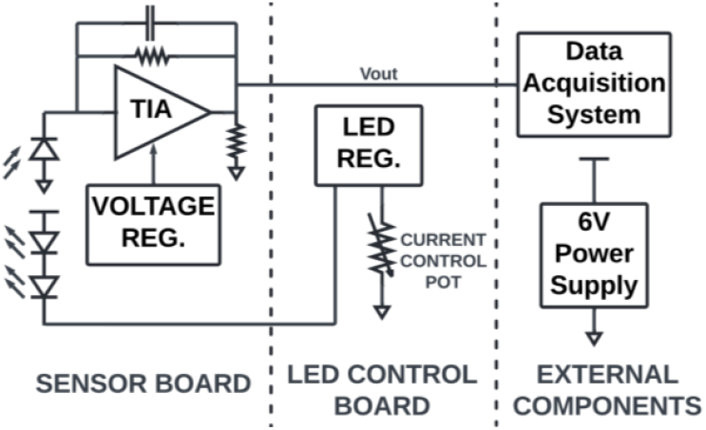
Schematic of the custom PPG sensor. The full setup consists of a custom flexible printed circuit boards (the sensor board) and a rigid printed circuit board (LED control board) while data is analyzed and extracted on a data acquisition system. “TIA” stands for Transimpedance Amplifier and “Reg.” stands for regulator

The sensor board had six LED slots which allowed for variable PD-LED spacing of 5, 12, or 20 mm (Fig. 3). To account for potential reductions in light received at the detector, these boards were designed to have a gain of 102 dB, 110 dB, and 120 dB at the transimpedance amplifier. After assembly, the system components were coated with a high-impact resistant, non-sagging epoxy for disposable medical devices (Loctite Hysol M-121HP Medical Device Epoxy Adhesive, Henkel Adhesive Technologies). Epoxy was only applied to solder joints, to waterproof and protect them during bending, and to prevent IR absorption on optoelectronic components.

**Figure 3.**
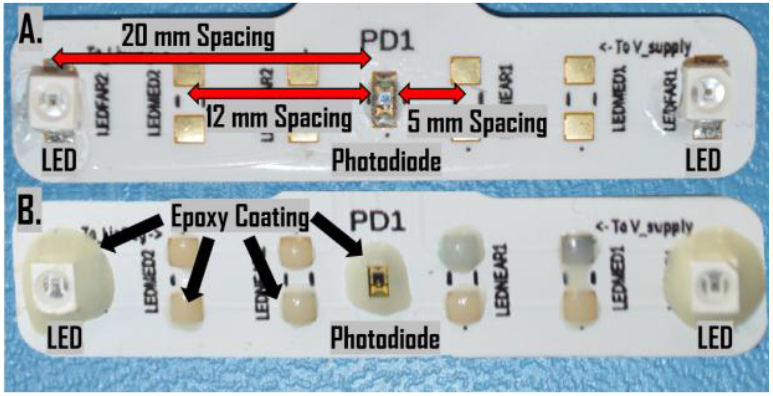
Top-down view of custom sensor boards. A show the front LED side of the printed circuit board prior to epoxy coating. B shows the same board after being coated by a medical grade epoxy.

A simple ray tracing model (assuming no scattering or absorption) predicted that farther spacing between the LED and PD would prevent reflection from tissue near the sensor but would lead to a mild focusing effect where light would reflect back only at deeper tissue depths, depending on the LED focal cone and PD capture angle. Additionally, a gradual reduction in PPG signal amplitude was expected due to continuous absorption of IR light by all tissue encountered (not just blood). Therefore, we expected that the highest amplitude signal will be reflected from the nearest LED spacing, which will quickly reduce as that target tissue becomes deeper. We also predicted that farther spacings of LED to PD would have a lower signal to noise ratio (SNR), but that an optimal reflection depth would exist for each LED-PD spacing. Using this model, we calculate the optimal vessel depth for a 5, 12, and 20 mm spacing as 2.3, 5.8, and 17.0 mm, respectively.

### B. Pulsatile Phantom

*In vitro* bench testing used an optical phantom of pulsatile blood flow to mimic the implant environment in humans. A few phantoms developed recently focused on the conditions in which PPG interacts with tissue, [10] however these phantoms did not emulate human hemodynamics of large vessels deep beneath the skin surface. A custom pulsatile flow phantom was developed to provide a repeatable *in vitro* testing environment to evaluate PPG performance. The bench model included two pumps, silicone tubing, and a data acquisition (DAQ) system (Fig. 4).

**Figure 4.**
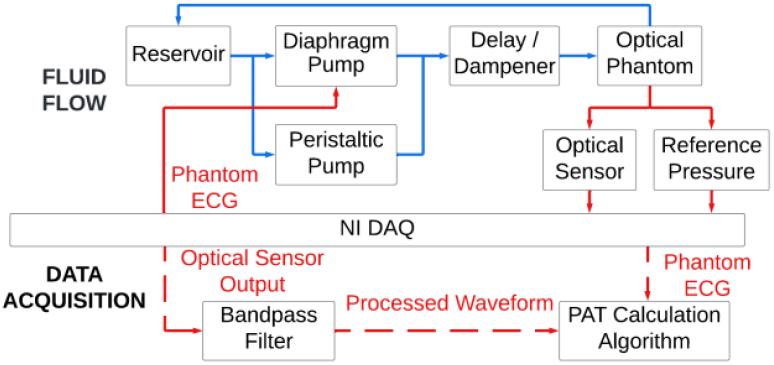
Block diagram of pulsatile phantom, optical phantom, and data processing. Blue lines indicate fluid flow, solid red indicates analog signal, dashed red indicates digital.

The pulsatile phantom used a blood-mimicking aqueous solution of 40% glycerol to mimic the viscosity and fluid dynamics of blood [11]. The fluid flowed through two parallel pumps: a diaphragm pump (bayite 12V DC Fresh Water Pressure Diaphragm Pump BYT-7A102) and a peristaltic pump (Cole-Parmer Masterflex L/S Peristaltic Pump - 07522-20). The peristaltic pump ran continuously to control the mean arterial pressure (MAP) of the system. The pulsatile pressure was generated by a National Instruments (NI DAQ) 5V, 10% duty cycle square wave output to trigger a DC solid state relay. The relay switched power to the diaphragm pump at 1 Hz with a pulsatile pressure swing of roughly 40 mmHg. Peak pulsation pressure was controlled by the diaphragm pump voltage provided from a lab power supply, which was varied between 4 and 7 V.

To better simulate human arterial hemodynamics, the solution passed through a dampener which included an isolated compressible air pocket. The solution then passed through a 2.4-m length of additional tubing to further smooth the arterial pressure waveform. An optical phantom was then inserted inline to the flow system. This optical phantom (described in the next section) mimicked the IR properties of tissue while enabling flow through a simulated embedded vessel. A luer-lock pressure sensor (PendoTECH MEMS-HAP™) was used to collect reference arterial pressure waveforms from the pulsatile phantom.

### C. Optical Phantom

The optical phantom was constructed from silicone rubber (Ecoflex™ 00-10, Smooth-On, Inc.). To mimic the IR absorption of human tissue, a dispersion of 0.6 mg/ml of India ink and a 2 mg/ml concentration of titanium dioxide (TiO2) [12], [13] was added to the silicone. The dispersion was created by suspending the additives in alcohol using ultrasonication for 10 minutes (Qsonica Q500), then mixing with the silicone materials. The IR-absorbing silicone was cast into a form around a 9.5 mm outer diameter silicone tube with a 1.5 mm thickness, which represented a blood vessel embedded in tissue. After curing the phantom for 12 hours at 22° C, finished phantoms were connected to the pulsatile system for optical signal validation (Fig. 6).

Three prototype PPG boards with LED to PD spacing of 5, 12, and 20 mm were tested (Fig. 5). PPG was recorded synchronously with reference pressure and the pulsatile pump control signal using the NI DAQ (Fig. 4). The pump control signal was recorded as an approximation to the electrocardiogram (ECG), because positive-going pulses in pump control preceded the arrival of a new pulse of blood-mimicking fluid, like the use of ECG recording in PPG PAT calculation. All data were further processed post-hoc in MATLAB R2023b.

**Figure 5.**
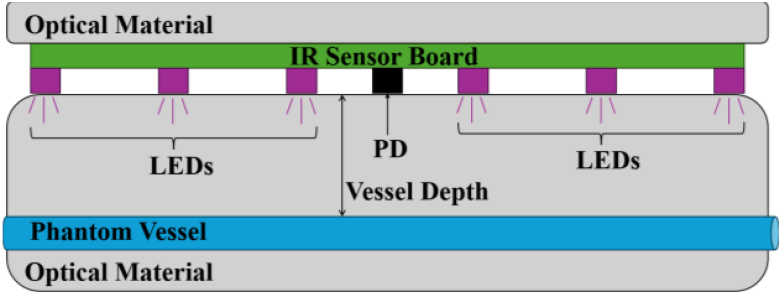
The optical phantom is constructed of EcoFlex silicone rubber, TiO_2_, and India ink. The PPG sensor was placed atop the phantom and a block of optical material simulated the subcutaneous environment.

**Figure 6.**
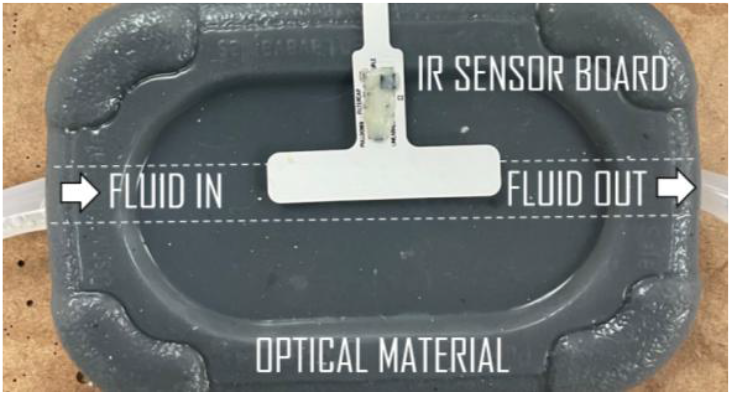
Top-down view of optical phantom. The dashed line represents the silicone vessel embedded in the block of optical material.

### D. In Vitro Experiments

To evaluate the signal quality, SNR was calculated by dividing the root mean square of the sensor waveform by the root mean square of the noise. The noise is recorded by taking the waveform recording of the device on the optical phantom with no activity from the pulsatile phantom. Two experiments are then done to visualize the device performance. In the first experiment, the sensor was placed directly over the location of the vessel and pressure is gradually incremented. The SNR is then calculated for each spacing and vessel depth orientation as well as its corresponding pressure in the vessel. In the second experiment, pressure was maintained at 126/78 mmHg and the sensor was shifted laterally away from the vessel in increments of 10 mm. SNR was then calculated for every vessel depth, LED spacing, and sensor placement to visualize how the signal quality changed across different orientations. An analysis of variance test (ANOVA) paired with a Tukey’s post-hoc test using a 95% confidence interval was done for each experiment to explore the difference between different conditions on the device performance.

### E. Data Processing

The PPG optical waveform and the 1 Hz pulsatile pump control waveform (treated as a phantom generated ECG) were analyzed. Data were filtered through a 12th-order Butterworth bandpass filter with 0.3 to 10 Hz passband. Windows were selected over each ECG period, and the first and second gradient of the optical waveform were analyzed for feature extraction [14]. This resulted in six features which were analyzed in parallel: maximum first gradient value, minimum first gradient value, zero first gradient value, maximum second gradient value, minimum second gradient value, and zero second gradient value [15]. Finally, the time delay was measured between each relevant feature and the beginning of the ECG window, and recorded as a finalized PAT.

### F. Preliminary In Vivo Experiment

A single acute non-survival experiment was conducted on an adult New Zealand White rabbit (4.1 kg body weight). Protocols involving animal experiments were approved by the Institutional Animal Care and Use Committee (IACUC) of Case Western Reserve University and with veterinarian oversight. The animal was sedated using inhaled isoflurane gas and mechanically ventilated. Vital signs were monitored continuously, and core temperature was maintained during anesthesia. Muscle fascia of the biceps femoris was exposed via a single long incision through the skin, to facilitate placement of the PPG sensor between the skin and muscle.

During PPG recording, blood pressure was maintained at 20 mmHg via intravenous fluid delivery. This pressure was much lower than in humans, so a smaller PPG signal was recorded. Further, to accommodate the thinner tissue in the animal (as compared to a human muscle), a PPG board with a 20 mm LED spacing was used (Fig. 7). ECG and PPG waveforms were recorded simultaneously. ECG was recorded with stainless steel electrodes (Rhythmlink 13 mm Subdermal Needle Electrode) and a CED 1902 isolated amplifier. Reference arterial pressure was measured by surgically inserting a micro-tip catheter (Scisense Catheter line, Transonic) into the femoral artery. While recording, a blood pressure rise was induced via intravenous phenylephrine (2 mg/ml, 3000 µl/min) to raise the MAP. Continuous measurements of PPG, ECG, and blood pressure were captured, and stored for post-hoc estimation of blood pressure via pulse arrival time (PAT) calculation.

**Figure 7.**
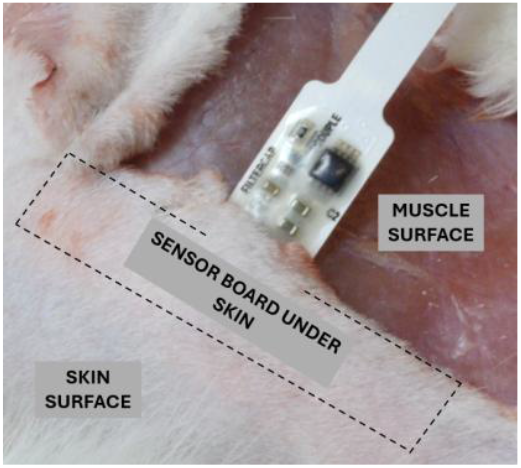
Image of optical sensor board placed subcutaneously on an animal. The dashed line represents the outline of the sensor board that is placed under the skin with the LED side facing the muscle surface.

## III. Results

### A. Pulsatile Waveform

Reference pressure measured in the optical phantom indicated a 40-mmHg pulsatile waveform was obtained over a range of baseline pressures. The mean arterial pressure (MAP) across all peristaltic pump settings ranged from 17 mmHg to 240 mmHg. For every four-minute window the waveform was analyzed and features were marked across each waveform without fail. The features 1-6 within an ECG peak time-window for a sensor board output are as follows: the maximum first gradient point, the minimum first gradient point, the zero first gradient point between the features 1 and 2, the maximum second gradient point, the minimum second gradient point, and the zero second gradient point between features 4 and 5 (Fig. 8).

**Figure 8.**
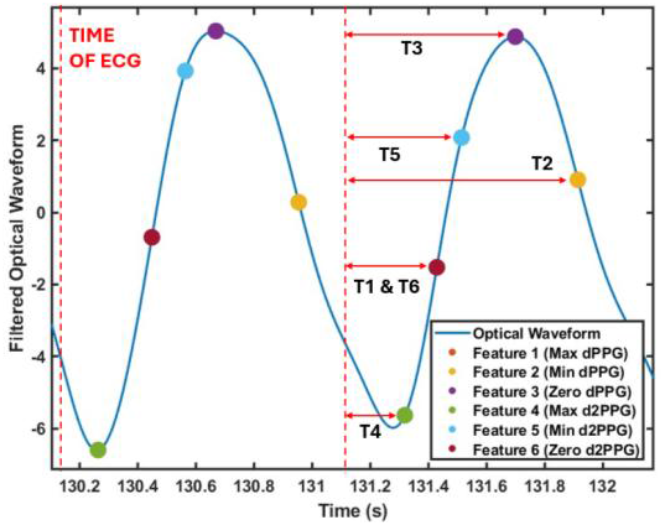
Filtered optical waveform, with marked features to measure the time delay between the phantom ECG and the select feature.

PAT could be calculated on the phantom at a rate similar to that of a human [16] as for feature 4 the phantom showed a 180 ms delay for an average healthy blood pressure of 126/78 mmHg. However, the expected inverse relationship between PAT and MAP reported in physiological literature was not observed from our phantom PPG [3].

### B. Signal Quality Evaluation

An analysis of variance (ANOVA) paired with a Tukey’s honestly significant difference test using a 95% confidence interval was successfully performed (Table 1) for the first *in vitro* experiment and showed some noticeable differences between the boards. The 5 mm spacing board showed a higher mean SNR than both the 12 mm and 20 mm board with a difference of 5.8 dB and 5.7 dB respectively. However, the test did not show any significant difference in mean SNR between the 12 mm and 20 mm spacing boards. Additionally, the 5 mm board showed a 13.9 dB reduction in mean SNR for vessel depth change from 10 mm to 20 mm. The 12 mm board showed an SNR reduction of 6.4 dB and 3.2 dB for a vessel depth change from 5 mm to 20 mm and from 15 mm to 20 mm. However, the 20 mm board showed no significant difference across different vessel depth despite visually showing a slight increase in SNR at 15 mm in Fig. 9.

**TABLE I.** Analysis of Variance Summary.

| Experiment 1 | ANOVA Summary |  |  |
| --- | --- | --- | --- |
|  | Degree of Freedom | F-Value | P-Value |
| Spacing | 2 | 8.42 | 0.001 |
| Vessel Depth | 3 | 4.61 | 0.005 |
| Experiment 2 |  |  |  |
| Spacing | 2 | 1.27 | 0.29 |
| Vessel Depth | 3 | 5.55 | 0.002 |
| Sensor Placement | 4 | 8.10 | 4.1e-05 |

**Figure 9.**
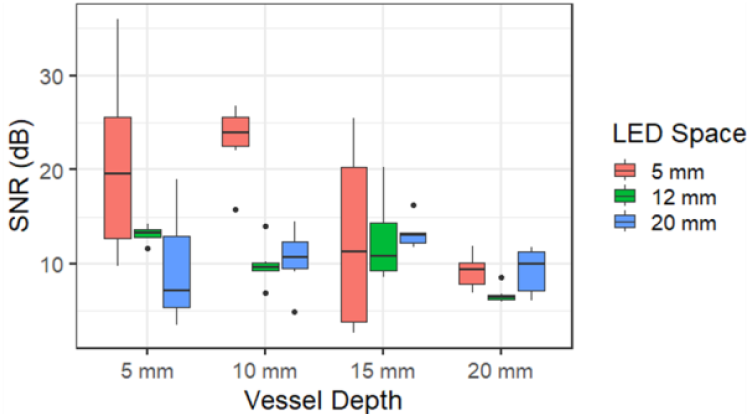
SNR of the PPG waveform was measured for each LED spacing and depths of 5 – 20 mm.

An equivalent variance test for the second *in vitro* experiment showed that for all spacings and positions, increasing vessel depth from 5 mm to 15 mm and 20 mm yielded a significant average SNR reduction of 5.8 dB and 7.9 dB respectively. While all other increases in vessel depth support an SNR reduction as the vessel depth increased, the 95% confidence interval showed potential for the reverse to be true. Additionally, for all vessel depths and spacings, there is a reduction in average SNR when the placement shifts from 0 mm to −20 mm (−8.8 dB), from 0 mm to +20 mm (−7.7 dB), and from 0 mm to −10 mm (−7.4 dB).

### C. Acute, Implanted PPG Experiment Results

The implanted PPG waveform demonstrated the same frequency to the ECG when observed intra-operatively and the signal quality was sufficient to enable real-time changes in amplitude with position changes. The sensor, however, was not overly sensitive to position and began producing a reliable and stable PPG signal as soon as it was placed on the biceps femoris. The sensor output displayed an SNR of 22.4 dB.

While infusing phenylephrine, MAP rose from 55 mmHg to 115 mmHg over a 250 second period. PPG-derived PAT measurement over the same period showed an inverse relationship (Fig. 11).

**Figure 10.**
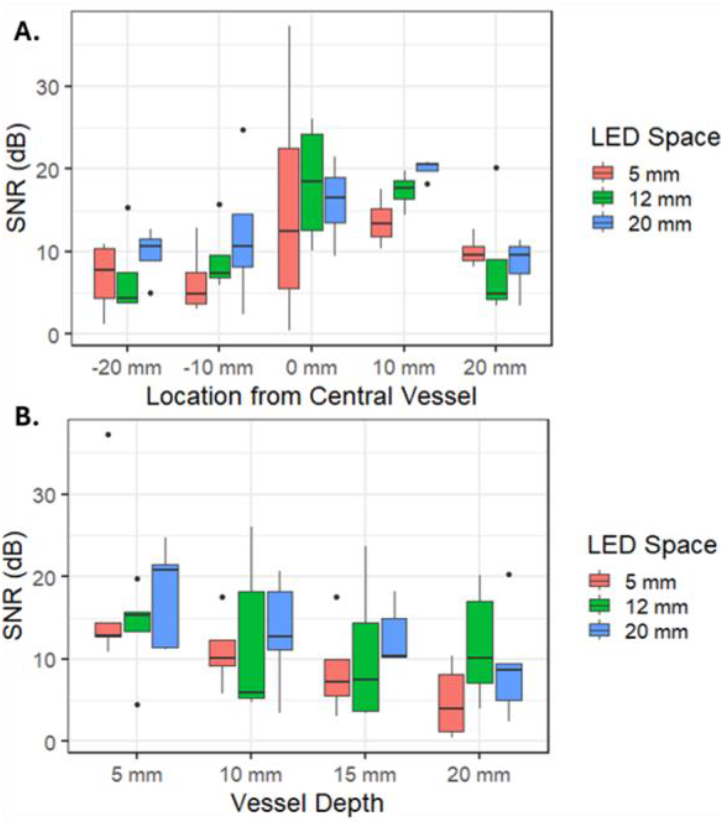
PPG SNR plot for different vessel depths and a lateral displacement between the sensor and vessel. (a) The best SNR was achieved directly above the vessel, regardless of LED spacing. (b) SNR dropped quickest with vessel depth for the 5 mm LED spacing.

**Figure 11.**
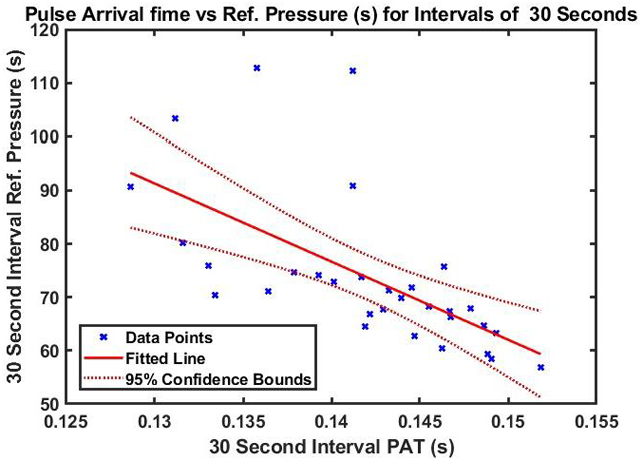
PAT results calculated from peak of R wave on ECG to Feature 4 (Maximum second gradient) of the filtered optical waveform. The x-axis represents the average PAT calculated within a 30 second interval for the animal trial. The y-axis represents the mean arterial pressure for that same interval. Results are showing an inverse relationship between the two variables with a relatively weak fit.

## IV. Discussion

The SNR figures and variance analysis suggest that for an increasing vessel depth, there is a detectible reduction in SNR within a 95% confidence interval. This reduction is likely due to the phenomenon of a deeper vessel requiring a longer path length for light to travel to reach the vessel and reflect to the detector. This behavior also matches what has been found on other pulsatile phantoms [17].

Additionally, the figures suggested that variations in the placement of the board away from the central vessel also result in a reduction of SNR. A potential explanation for reduced SNR is that shifting the sensor board away from the central vessel reduces the amount of light emitted by the LED, which has a limited viewing angle, from to reaching the vessel. Finally, we found that holding the sensor board’s position constant over the vessel while the pressure is varied, increasing vessel depth reduced SNR for the 5 mm spacing board and 12 mm spacing board, but not for the 20 mm spacing board. Figure 9 suggested that the 20 mm spacing had peak average SNR at the 15 mm vessel depth, which is consistent with our prediction that the optimal vessel depth would be 17 mm, but the variance test cannot currently confirm this. However, the inability to find a significant reduction in SNR for the 20 mm board could suggest a potential benefit in choosing a farther spacing orientation for a subcutaneous sensor. If a farther spacing reduces the variance in SNR for varying optical conditions, it could potentially provide more reliable acquisition of a PPG signal regardless of the environment. However, future experimentation is needed to confirm whether this hypothesis is valid. Additionally, it is noted that the environment that the phantom hopes to mimic is a muscle surface with embedded capillaries. Since the phantom consists of a single vessel, it is likely to better replicate a single, large, arterial vessel instead of a bed of capillaries that the sensor would be subject to when placed on the muscle surface.

The inverse relationship with MAP and PAT obtained *in vivo* during phenylephrine infusion matched prior reports [3], [18] suggesting that PAT decreases as blood pressure increases, due to increased vascular rigidity leading to increased pulse wave velocity in arteries. In this experiment, however, a low correlation of r^2^ = 0.385 was obtained. We hypothesize that variations in the feature detection algorithm and a limited recording time for a singular acute experiment may have resulted in this low correlation. These aspects may be improved via signal processing, optoelectronic improvements in the sensor, and increased animal recording times.

## V. Conclusion

In this paper a custom PPG setup and phantom was created to explore the potential of a subcutaneous setup with varying spacing between the light source-to-detector spacings as a method to reduce reliance of signal quality on capillary density. In this investigation there is an observed trend of increased SNR for deeper vessel depths for the farthest LED to detector spacing which is aligned with our initial prediction. However, ANOVA and post-hoc tests reveal that these trends did not reach statistical significance. While these results are encouraging towards improving the reliability of signal acquisition for subcutaneous use, further investigation is needed with improvements towards phantom construction, sensor construction, and statistical power to validate this phenomenon.

